# A single cycle influenza virus coated in H7 hemagglutinin provides heterotypic protection and neutralising antibody responses to both glycoproteins

**DOI:** 10.1101/224550

**Authors:** Timothy J Powell, Pramila Rijal, Rosanna M McEwen-Smith, Haewon Byun, Marc Hardwick, Lisa M Schimanski, Kuan-Ying A Huang, Rodney S Daniels, Alain R M Townsend

**Author notes:** Current address: Respiratory Medicine Unit, NIHR Biomedical Research Unit, Nuffield Dept of Clinical Medicine, University of Oxford, John Radcliffe Hospital, Headley Way, Oxford, OX3 9DU, UK. To whom correspondence should be addressed Prof Alain Townsend: ^a^MRC Human Immunology Unit, MRC Weatherall Institute of Molecular Medicine, University of Oxford, John Radcliffe Hospital, Oxford, OX3 9DS, UK.

## Abstract

A non-replicating form of pseudotyped influenza virus, inactivated by suppression of the hemagglutinin signal sequence (S-FLU), can act as a broadly protective vaccine. S-FLU can infect for a single round only, and induces heterotypic protection predominantly through activation of cross-reactive T cells in the lung. Unlike the licensed live attenuated virus, it cannot reassort a pandemic HA into seasonal influenza. Here we present data on four new forms of S-FLU coated with the H7 hemagglutinins from A/Anhui/1/2013 and A/Shanghai/1/2013, H7N9 viruses that emerged recently in China, and A/Netherlands/219/2003 and A/New York/107/2003. We show that vaccination in the lung induced a strong local CD8 T cell response and protected against heterosubtypic X31 (H3N2) virus and highly virulent PR8 (H1N1), but not influenza B virus. Lung vaccination also induced a strong neutralising antibody response to the encoded neuraminidase. If given at higher dose in the periphery, H7 S-FLU induced a specific neutralising antibody response to H7 hemagglutinin coating the particle. Polyvalent vaccination with mixed H7 S-FLU induced a broadly neutralising antibody response to all four H7 strains. S-FLU is a versatile vaccine candidate that could be rapidly mobilized ahead of a new pandemic threat.

## Introduction

Influenza virus causes annual widespread respiratory tract infections resulting in mortality and morbidity in the human population. They can be divided into subtypes and genetic groups according to the hemagglutinin (HA) and neuraminidase (NA) glycoproteins on their surface [1]. Influenza can generate novel virus variants by genetic reassortment during co-infection with two or more strains, and these can spread rapidly through an unexposed human population to cause pandemics. Recent zoonoses of novel avian influenza viruses, such as H7N7 and H9N2, were localized and caused mild disease [2, 3]. In contrast a novel virulent strain of influenza H7N9 (group 2), not previously isolated in humans, appeared in China from March to May 2013 [4]. Since then there have been 1564 confirmed human infections with influenza H7N9 with at least 612 deaths including an unexpected increase in cases during the 2016/7 influenza season (WHO Risk Assessment 27 September 2017). To date, this virus has been weakly transmissible between humans, but it is concerning that it may need to make only a small number of amino acid substitutions in HA to enable human-to-human transmission [5], and humans may not have any neutralising antibody. By contrast, during the 2009 H1N1 pandemic it was seen that older people had some protection from infection through cross-reactive antibody induced by earlier H1N1 virus infections [6–8]. New pandemics can originate from either group 1 or 2 influenza A viruses, so cross protection between genetic groups has recently been defined as a major goal for developing broadly protective influenza vaccines [9].

Recovery from influenza infection can provide immunity that is cross protective between A viruses in the absence of detectable neutralising antibodies. This was first noticed in 1933 through comparison of infections with swine and human influenza (distantly related H1N1 viruses) in ferrets [10]. It was named “partial specific heterotypic immunity” by Schulman and Kilbourne [11] in 1965, who observed partial protection against an H2N2 virus after recovery from an H1N1 infection in mice. It has been demonstrated in several species including mice, ferrets, guinea pigs, poultry, swine, and non-human primates (reviewed in [12, 13]), and is associated with reduced virus replication and pathology in the lung, and reduced transmission [14, 15]. Optimal induction of heterotypic immunity requires infection of the lung or total respiratory tract as opposed to infection at a peripheral site, including the nose [16–21].

In mice the cross protective effect of prior influenza infection is associated with a cross-reactive Cytotoxic T Lymphocyte (CTL) response, and can be transferred with specific class I restricted CD8 CTLs [22–26]. Both murine and human CD8 T cells recognize peptides derived from conserved virus proteins processed in the cytosol of infected cells [27–30]. While cell depletion and gene knockout experiments confirm a central role for T cells, there is evidence for additional contributions from B cells and innate immune cells in heterotypic immunity [31–33].

Heterotypic protection has been more difficult to demonstrate in humans (reviewed in [12, 13]), and may be limited in duration [34]. However, two recent observational studies correlated milder symptoms and reduced virus shedding with strong CD8 T cell memory during the recent H1N1 pandemic [35, 36]. Studies have also demonstrated cross-reactive T cells that recognize conserved components of the H7N9 and H5N1 viruses in individuals exposed only to seasonal influenza [37, 38], and a further study correlated high levels of cross-reactive CD8 T cells with milder disease in humans hospitalized with H7N9 infection [39]. These observational studies on natural influenza infections support evidence for T cell associated protection of humans from experimental influenza [40, 41].

The sum of evidence suggests that the protective mechanism of heterotypic immunity to influenza is multifactorial with CD8 T cells playing a central role. For this reason it seems logical that vaccines designed to induce it should be as closely related to live influenza as possible. We previously demonstrated that a novel replication deficient influenza virus vaccine, based on inactivation of the HA signal sequence ([42] hence “S-FLU”), coated with H1 hemagglutinins can induce cross protective immunity against PR8 (H1N1, group 1) and X31 (H3N2, group 2) viruses which have divergent surface glycoproteins but the same internal genes as S-FLU [43].

We have shown that in ferrets immunization with H1N1 or H5N1 S-FLU (group 1) significantly reduced replication of H1N1, H6N1, H5N1 (group 1) and H7N9 (group 2) wild-type viruses in the lung [44]. In pigs, immunization with H1N1 or H5N1 S-FLU reduced virus titres in nasal swabs and lungs following challenge with a swine H1N1pdm09 isolate and [45]. S-FLU can infect cells for one round only, but this is sufficient to express all the conserved virus proteins in the cytosol for induction of a vigorous local CD8 T cell response. Other similar non-replicating influenza constructs were also able to protect animals vaccinated in the lung (reviewed in [46]).

S-FLU viruses are similar to Live Attenuated Influenza Virus (LAIV) [47, 48] in terms of being able to stimulate strong T cell mucosal immunity and heterotypic protection in mice and ferrets [43, 46], but have two major advantages over the licensed form of LAIV. S-FLU is unable to replicate so vaccinees should not shed virus [49], and in addition S-FLU cannot donate a pandemic HA sequence to seasonal influenza by reassortment because it is coated in the HA glycoprotein, but does not contain the pandemic vRNA encoding HA [43]. We show here that vaccination in the lung with a novel form of S-FLU that is pseudotyped (coated) with H7 hemagglutinin from the A/Anhui/1/2013 H7N9 influenza virus protects mice against challenge with viruses of different HA subtype, highly lethal PR8 (H1N1 Cambridge strain) and X31 (H3N2), but carrying the same conserved internal genes as the S-FLU constructs. Protection after intranasal (i.n.) administration is associated with a strong local crossreactive CTL response in the lung.

Lung immunization with protective doses of S-FLU is not able to generate a reliable neutralising antibody response to the HA glycoprotein coating the pseudotyped particle [43], probably because the S-FLU infected cells cannot synthesize HA. However, we now demonstrate that the priming dose of S-FLU in the lung does induce a strain specific neutralising antibody response targeting the encoded NA, which unlike the HA is synthesized by infected cells. Further, H7-S-FLU given in a larger dose in the periphery (intraperitoneally (i.p.)) can generate a strong neutralising antibody response both to the H7 HA coating the particle, in addition to the encoded N1 NA. Administration of a mixture of S-FLU, coated in diverse H7 HAs, induced a broadly neutralising antibody response to the four viruses in the mix (including Eurasian and North American strains), as well as cross-reactive T cells in the spleen.

S-FLU, because it closely mimics natural influenza infection, but can neither replicate nor reassort an HA vRNA, may function as a versatile pre-pandemic vaccine for the induction of cross protective T cells to the conserved internal proteins and neutralising antibodies to the glycoproteins coating the virus.

## Materials and Methods

### Media, Reagents and Tissue Culture

Virus Growth Media (VGM) was DMEM with 0.1% bovine albumin (Sigma A0336), 10 mM HEPES, 2 mM glutamine, 100 U/mL penicillin and 100 μg/mL streptomycin (all from Sigma, Poole UK). D10 was DMEM with 10% v/v foetal calf serum (Sigma) plus glutamine and antibiotics as above. R10 was RPMI (Sigma) with 10% serum, glutamine and antibiotics. MDCK-SIAT1 cells [50] were obtained from the European Collection of Cell Cultures (ECACC) distributed by Sigma.

### Influenza Viruses

Plasmids carrying the genes of the Cambridge strain of A/Puerto Rico/8/34 were used to generate PR8 virus [43]. A reassortant containing the HA and NA from A/Aichi/2/68 (H3N2) and internal genes of PR8, X31[51], was a gift from J McCauley (Crick Worldwide Influenza Centre, The Francis Crick Institute, London). Viruses were produced as culture supernatants from MDCK-SIAT1 cells [50] using standard methods and were stored at −80°C after centrifugation.

### Antibodies to Influenza

Human monoclonal antibodies (mAbs): Anti-NP 2-8C has been described [43]. 9B-26 specific to H7 HA from A/Anhui/1/2013, was isolated from a plasmablast FACS sorted from a peripheral blood sample provided by a donor post recovery from infection using standard methods [8, 52], and was screened for binding to MDCK-SIAT1 cells transduced to express H7 protein from A/Anhui/1/2013 (described below). Hyper-immune sheep sera: raised against HAs from A/New York/107/2003 (H7) (Cat No 09/148), A/mallard/Netherlands/12/2000 (H7) (Cat No 07/278) and A/VietNam/1194/2004 (H5) (Cat No 07/148), together with antisera raised against NA from inactivated reassortant H7N1 virus carrying N1 from A/California/07/2009 (H1N1pdm09) (Cat No 10/218), were obtained from the National Institute for Biological Standards and Control (Potters Bar, UK). Murine mAbs: NA2-1C1 anti-NA [53] and H9D3.Cb13 anti H1 from A/PR/8/34 were gifts from Jonathan Yewdell. CD6 to N1 from A/California/07/2009 [54], was a gift from Maryna Eichelberger;

### Nomenclature for S-FLU

To distinguish the various combinations of genotype and surface HA of the pseudotyped S-FLU vaccine viruses, we refer to the HA and NA genotypes between square brackets, followed by the origin of the surface HA derived from the cell line in which the virus was propagated. The S-FLU viruses we have produced have a near full length HA vRNA derived from A/PR/8/34 that has the signal sequence inactivated to prevent surface expression [42] and in some the HA coding sequence has been replaced with eGFP to provide a marker in infected cells.

The pseudotyped viruses produced for this paper were formally [S-eGFP/N1(PR8)]. H7 (Anhui/1/2013); [S-eGFP/N1(PR8)]. H7 (Shanghai/1/2013); [S-eGFP/N1(PR8)]. H7 (New York/107/2003); [S-eGFP/N1(PR8)]. H7*t (Netherlands/219/2003) with the polybasic cleavage site replaced with a trypsin site; [S-eGFP/N1(PR8)]. H5*t (Vietnam/1203/2004) with the polybasic cleavage site replaced with a trypsin site; [S-eGFP/N1(Eng195)]. H1/G222D/K154E (England/195/2009) with the amino acid substitutions at the positions indicated to improve virus yield [55].

### Production of S-FLU

The development of S-FLU has been described in detail [43]. The S-eGFP vRNA that replaces the HA vRNA was derived by suppression of the PR8 HA signal sequence and replacement of the coding region with eGFP sequence and additional safety mutations. The N1 neuraminidase, and all the internal genes, for the H7 S-FLU viruses were derived from PR8. Novel S-FLU pseudotype viruses were produced by propagating an S-FLU produced earlier in a panel of MDCK-SIAT1 cells stably transduced with the lentiviral vector pHR-SIN [56] engineered to express full-length codon optimised HAs, but lacking 3’ and 5’ UT regions, from a variety of viruses: H7 A/Anhui/1/2013 (GISAID EPI439507), H7 A/Shanghai/1/2013 (GISAID EPI439486), H7*t A/Netherlands/219/2003 (Genbank AY338459.1, with the polybasic cleavage site replaced with the trypsin sensitive site from A/Anhui/1/2013), H7 A/New York/107/2003 (Genbank EU587368.1), H5*t A/Viet Nam/1203/2004 (GenBank: EF541403.1 with polybasic cleavage replaced with a trypsin site), H1/G222D/K154E A/England/195/2009 (Genbank ACR15621.1 carrying the amino acid substitutions indicated to enhance virus yield [55]. Transduced cells were stained with H7, H5 or H1 HA-specific human mAbs (prepared in house) and FACS sorted to achieve maximal expression of HA.

To produce new versions of S-FLU encoding either N1 (PR8) or pdm09N1 (A/Eng/195/2009), and coated with various HAs, these transduced MDCK cells were seeded with S-FLU, either [S-eGFP/N1(PR8)].H1(PR8) or [S-eGFP/N1(PR8)].H5*t (VietNam/1203/2004) or [S-eGFP/N1(Eng195)].H1(Eng195) [43] at multiplicities of infection (MOI) in the range 0.01-1 in VGM. After 1 hr cells were washed and cultured in VGM containing TPCK trypsin (Sigma T-1426) at 0.5–1 μg/mL and virus was harvested after 48 hrs and stored in aliquots at −80°C.

### In Vitro Characterization of S-FLU viruses

Virus titration: NP expression within infected cells was detected after overnight culture of MDCK-SIAT1 cells incubated with titrated doses of virus. NP protein was detected using human anti-NP mAb (2-8C, produced in house) on permeabilised cells followed by anti-human IgG-HRP and then TMB POD substrate (Roche, Welwyn Garden City) which after addition of 1M H_2_SO_4_ was read at 450 nm on a plate reader (Bio-Rad). Viruses expressing eGFP were assessed by infection of MDCK-SIAT1 cells with eGFP detection in a CLARIOstar fluorescence plate reader as described by Martinez-Sobrido et al., [57]. The vRNAs of ‘Bright’ viruses that expressed high levels of eGFP, cloned by limiting dilution in MDCK-SIAT1 cells, were found to have a single base mutation of T60C, compared to their ‘Dull’ counterparts. This mutation destroyed an artefactual translation initiation site upstream of the eGFP translational start site at base 96. Titration mid-points for NP expression and eGFP fluorescence were equivalent.

### Virus Quantification

The 50% Tissue Culture Infectious Dose (TCID_50_) was determined in MDCK-SIAT1 and H7 transfected MDCK-SIAT1 cells using monolayers set down after overnight culture in D10. Cells were washed with PBS and infected with a dilution series of viruses in VGM for at least 30 min, then 150μL of TPCK trypsin containing VGM was added to give a final concentration of 1μg/mL and plates were incubated for a further 2-4 days at 37°C. TCID_50_ was calculated using the Reed & Muench method of cumulative percentage of positive and negative wells [58]. Virus in the supernatant was quantified by either hemagglutination of 1% human red blood cells (RBCs), or by NP staining or eGFP detection in infected MDCK-SIAT1 monolayers. Typical HA and TCID_50_ titres of the H7 S-FLU viruses were A/Anhui/1/2013: 16 HAU/50μl (1.5 × 10^7^ TCID_50_/ml), A/Shanghai/1/2013: 4 HAU/50μl, A/Netherlands/219/2003: 128 HAU/50μl (3.3 × 10^8^ TCID_50_/ml), A/New York/107/2003: 64 HAU/50μl. The H7 S-FLU expanded in MDCK monolayers expressing A/Netherlands/219/2003*t consistently reached the highest titres.

### Microneutralisation (MN) Assays

Were performed as described [43] with minor modifications. The assay measures blockade of virus entry without trypsin present based on the method of Rowe et al., [59]. Briefly, viruses were titrated to give saturating infection of 3×10^4^ MDCK-SIAT1 cells/well in 96 well flat-bottomed plates as detected either by NP expression, or eGFP fluorescence [57]. Dilutions of antibody were mixed with virus and incubated for 1–2 hr at 37°C. Indicator MDCK-SIAT1 cells (3×10^4^/well) were added and the mixtures incubated overnight. The suppression of infection was measured either by reduced NP staining on fixed and detergent treated cells, or by fluorescence of fixed cells measured on a CLARIOstar fluorescence plate reader. Murine sera were heat inactivated for 30 min at 56°C. The pre-prepared sheep sera from NIBSC were used without additional heat inactivation.

### Enzyme Linked Lectin Assay (ELLA) for Neuraminidase Inhibiting Antibody

The assay was performed as described [60, 61] with minor modifications. The H7 S-FLU vaccines used to induce the NA response encoded the N1 neuraminidase from PR8. Sources of N1 neuraminidase for the assay were provided by versions of S-FLU selected to avoid interference by neutralising antibodies to HA: i) PR8 N1 NA was provided by [S-eGFP/N1(PR8)].H5*t(VN/1203/2004) or PR8 wild-type (wt) virus ii) pdm09N1 NA by [S-eGFP/N 1 (Eng)]. H1(Eng/D222G/K154E) and iii) N2 (A/Aichi/2/1968) NA by X31 virus [51].

Antisera were heat inactivated at 56°C for 30min and 75μL plated out in doubling dilutions in VGM. Sources of NA were added in an equal volume of VGM (at a dose titrated to give maximum digestion of fetuin) and the mix incubated for 30 min at room temperature. 100μL of the mixture were removed and added to ELISA plates coated in fetuin as described [60, 61] and incubated for 18hr at 37°C in a CO_2_ incubator (which buffers the VGM). The plates were washed ×4 with PBS, PNA-HRP (Sigma L7759, 1 μg/mL in PBS/0.1% BSA) added for 2hr, washed ×4 with PBS and developed with 50μL eBiosciences TMB (Thermo Fisher), or OPD (Sigma P8287) for 5-20 min, stopped with 50μL 1M H_2_SO_4_ and read at 450 nm (TMB) or 492 nm (OPD). Titres are expressed as IC_50_: the dilution of antiserum that reduced NA activity by 50%, calculated by linear interpolation between the two neighboring points on the linear part of the dose-response curve.

### Flow cytometry

Monolayers of MDCK-SIAT1 cells, 10^6^ cells per well in 6 well plates, were infected with 50 hemagglutinating units (HAU) of PR8, H7 S-FLU or mock infected. After overnight incubation in D10 the cells were trypsin EDTA treated (Sigma) and labeled for FACS using human anti-H7 antibody (9B-26) or anti-N1 (NA2-1C1) or anti-PR8 HA (H9D3.Cb13), followed by anti-human or anti-mouse IgG secondary antibody conjugated to AL647 (Invitrogen). Positive control MDCK-SIAT1 cells transfected with H7 were labeled with human anti-H7 antibody 9B-26 followed by anti-human IgG AL647 (Invitrogen).

Enumeration of antigen specific CD8 T cells in C57BL/6 mice was done on freshly isolated blood collected in Microvette CB 300 tubes (Sarstedt), after lysis of RBCs with ammonium chloride solution, and then labeled with anti-mouse TCR-FITC (H57-597), anti-CD8 PE-Cy7 (53-6.7), anti-CD19 V450 (eBio1D3) and NP366-74 tetramer produced in house (by JP Jukes and D Shepherd) and conjugated to streptavidin APC by D. Puleston [62]. Flow cytometric events were collected on a Dako Cyan cytometer (Dako) and analysed using FlowJo (Tree Star Software). Cells that did not express CD19 but were positive for TCR were assessed for CD8 positivity and the percentages of tetramer positive cells, within the CD8+ gate, were enumerated in graphs. In some experiments mice were killed, their spleens and lungs taken and converted to single cell suspensions in Petri dishes by teasing through nylon mesh. After RBC lysis, cells were treated for FACS staining as for fresh blood. Cells were counted by trypan blue dye exclusion to enable absolute numbers of viable tetramer positive cells to be calculated. Absolute numbers were calculated by multiplying counts by percent in lymphocyte gate × percent in CD19neg/TCRpos gate × percent within CD8pos/Tetpos gate.

### Vaccination of Mice

BALB/cOlahsd or C57BL/6 female mice were obtained from the University of Oxford BMS JR SPF facility or from Harlan (Shaw Farm, Bicester). Mice were immunized intranasally with 50 μL S-FLU virus (doses are shown in the Figures) under inhaled Isoflurane (IsoFlo, Abbott, UK) anesthesia, which exposes the whole respiratory tract to the inoculum [63], at day 0 and day 14-16. We refer to this as “lung immunization”. Animals were routinely monitored at least twice weekly by observation and weighing. Challenge with PR8 or X31 virus took place at least a further 4 weeks later and was done using intranasal challenge as above with virus diluted in VGM. Weight loss and gain was monitored daily and morbid animals were humanely killed by cervical dislocation or inhalation of carbon dioxide. Morbidity was judged using 20% weight loss as an end point or a clinical scoring system [64]. Conditional survival was the percentage of animals not reaching 20% weight loss or a threshold clinical score or a combination of both. During the experiments one vaccinated animal unexpectedly died after challenge having lost 5% body weight, and one animal mock vaccinated with Viral Growth medium died before challenge.

### Virus Isolation from Lungs

Virus concentrations from infected lungs were measured as described previously [43]. Briefly: lungs were removed from freshly killed mice, weighed and snap frozen in liquid nitrogen. Thawed lungs were homogenized using a QiaShredder (Qiagen, UK) and centrifuged at 12000 rpm for 1 min. Supernatants were recovered and virus titre measured using TCID_50_ as described above with the additional step of filtration of the diluted lung through a 0.22 μm spin filter (Millipore, UK) prior to plating.

### Antibody and T cell assays

Samples of mouse sera and immune organs were treated and analyzed as previously described [43] and were assayed for MN and ELISA activities using similar techniques apart from the fact that H7 S-FLU was used in MN assays in addition to PR8 and X31. ELISA assays used purified soluble H7 glycoprotein produced in stably transduced 293T cells as described [43]: plates were coated with a 20 μg/mL solution (in PBS) for 2 hours and blocked overnight with 5% milk. ELISPOT assays were performed as previously described [43] using NP_147_ and HA_519_ CD8+ T cell epitopes. The NP_147_ sequence is shared by the priming S-FLU and challenge viruses because they are all derived from PR8 but HA_519_ is in PR8 HA only.

### Histological analysis of lungs

Lungs were aseptically removed from animals after CO_2_ euthanasia and exsanguination. Lungs were then put into 10% formalin and processed by the Oxford Centre for Histology Research (OCHRe, John Radcliffe Hospital, Oxford) paraffin embedded and stained with hematoxylin and eosin.

### Statistical Analysis

Data were analysed using Graph Pad Prism (v6 Graphpad Software) or SPSS (IBM, Portsmouth, UK) using either ANOVA for three or more sets of values or Students unpaired two-tailed t test. Data sets that did not meet the criteria of normal distribution were evaluated using non-parametric comparisons. Correlation was assessed using a Spearman test. P values < 0.05 were considered significant.

### Ethics Statement

Human antibodies to influenza NP 2-8C [43] and H7 hemagglutinin (9B-26) were obtained from human subjects in compliance with good clinical practice guidelines and the Declaration of Helsinki. The protocol was approved by the Research and Ethics Committee of Chang Gung Memorial Hospital, Taiwan and the Weatherall Institute of Molecular Medicine, Oxford. All subjects provided written informed consent.

## RESULTS

### Growth Properties of H7 S-FLU

Initially we assessed whether H7 S-FLU infected susceptible cells and expressed NP protein at a similar level to PR8 virus. Viruses were titrated in 96 well plates and then MDCK-SIAT1 cells were placed into each well and incubated overnight. Following labeling with anti-NP mAb (2-8C) and secondary anti-human HRP, we showed that there was a similar level of NP expression for both H7 S-FLU and PR8 virus (Figure 1A). We then added titrated viruses to MDCK-SIAT1 monolayers in the presence of trypsin and showed that only H7 (Anhui/1/2013) transfected MDCK-SIAT1 cells supported replication of H7 (Anhui/1/2013) S-FLU virus (Figure 1B, C): it replicated to titres of 1-2 ×10^7^ TCID_50_/ml. By contrast S-FLU seeded onto MDCK-SIAT1 cells expressing H7/Netherlands/219/2003 consistently achieved ~ 20 fold higher titres ~3 ×10^8^ TCID_50_/ml, similar to the PR8 control which reached ~2 × 10^8^ TCID_50_/ml (not shown).

**Figure 1.**
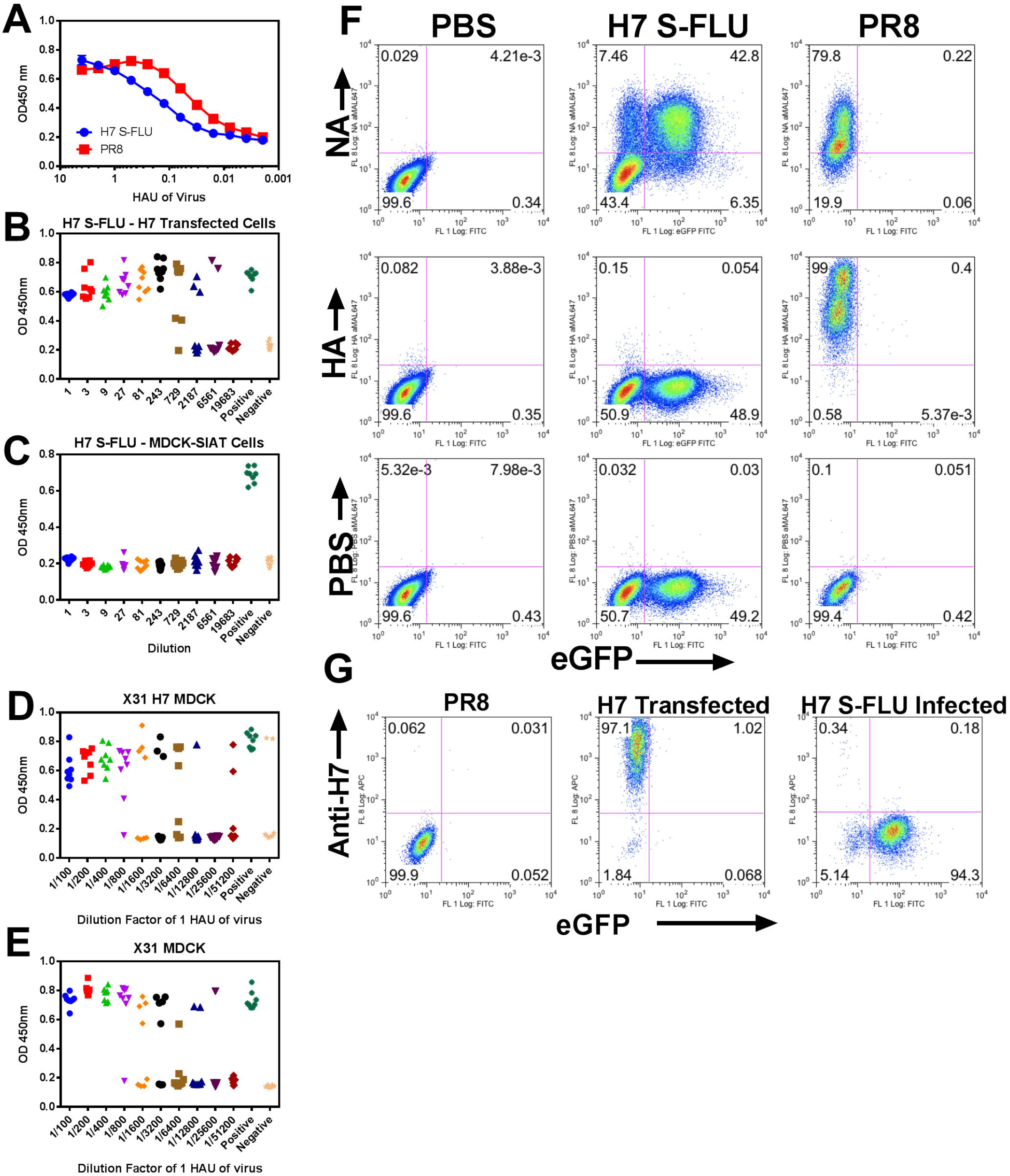
H7 S-FLU can infect MDCK-SIAT I cells but is unable to replicate without H7 HA expression in MDCK-SIAT1 cells. (A) MDCK-SIAT1 cells were added to wells containing titrated amounts of PR8 virus or H7 S-FLU and then incubated overnight. NP expression in cell monolayers was assessed by labeling with human anti-NP and anti-human IgG conjugated to HRP followed by color development. Values are means of two replicates with SD shown. Viruses were titrated onto (B) H7 (Anhui/1/20130 transduced MDCK-SIAT1 or (C) MDCK-SIAT1 monolayers and after three days in the presence of trypsin levels of NP were measured by ELISA. (D,E) X31 replicates in H7 transfected and WT MDCK-SIAT1. Titrated doses of X31 virus, in the presence of TPCK trypsin, were added to cell monolayers of MDCK-SIAT I transfected with H7 (D) or MDCK-SIAT1 (E) in 96 well plates. Three days later NP expression was determined by ELISA. (B, C, D, E) Eight replicates are shown for each dilution. (F) MDCK-SIAT1 cells were infected overnight with 50 HAU of H7 S-FLU or PR8 virus and then labeled for flow cytometry the next day. (G) MDCK-SIAT1 infected with H7 (Anhui/1/2013) S-FLU or mock infected, or MDCK-SIAT1 expressing H7 glycoprotein from a transduced cDNA were labeled with anti-H7 human mAb 9B-26: results of a representative experiment repeated at least twice are shown.

X31 virus grew to similar titres in both H7 transfected MDCK-SIAT1 and MDCK-SIAT1 cells implying that the transfection of MDCK-SIAT1 cells with H7 HA did not interfere with their ability to support virus replication (Figure 1D, E). Thus as shown with PR8 H1 S-FLU and Eng195 H1 S-FLU, the H7 S-FLU required a source of HA glycoprotein for productive virus replication.

To show that cells infected with H7 S-FLU were unable to express H7 HA, cells were mock infected overnight with PBS or infected with A/Anhui H7 S-FLU or PR8 wt virus and then labeled for FACS analysis the following day. H7 S-FLU infected cells expressed the eGFP protein as well as the NA from PR8 with the level of NA expression being similar to cells infected with PR8 wt (Figure 1F, top row). A small proportion of infected cells appeared to express NA without eGFP. This may be similar to the phenomenon described by Brooke et al., [53], where a proportion of influenza viruses fail to express protein from at least one vRNA in infected cells in vitro. Controls using anti-PR8 HA (Figure 1F middle row) showed that only PR8 infected cells had PR8 HA on their surface. Next, a human anti-H7 mAb 9B-26, developed in our laboratory, was used to stain cells. H7 transfected MDCK-SIAT1 cells expressed a high level of H7 HA (Figure 1G middle panel) while, by comparison, the H7 S-FLU infected MDCK-SIAT1 cells did not express H7 HA as demonstrated by lack of binding of human anti-H7 mAb to H7 S-FLU infected MDCK-SIAT1 cells (Figure 1G, right panel).

### T cell and antibody generation after vaccination in Lung

#### T cells

Vaccination or challenge was done by instilling 50μL of virus to the nose of an anesthetized mouse. Under these conditions at least 50% of the inoculum reaches the lung [63]. To determine the kinetics of T cell priming after lung vaccination with one or two doses of H7 S-FLU we instilled C57BL/6 mice with a dose of 16 HAU (7×10^5^ TCID_50_) H7 (Anhui/1/2013) S-FLU and then monitored generation of NP 366-74 specific T cells. This showed that the H7 S-FLU induced a population of CD8+ T cells that were present in the blood after one dose, increased after a second dose, and remained in circulation for at least 63 days (Figure 2A). On day 64 we killed groups of four mice and determined the percentages of tetramer positive cells that persisted in lung and spleen (Figure 2B). At day 64 there was a population of ~6% CD8+ tetramer binding cells present in the lungs of immunized animals that was not present in VGM primed mice (Figure 2C).

**Figure 2.**
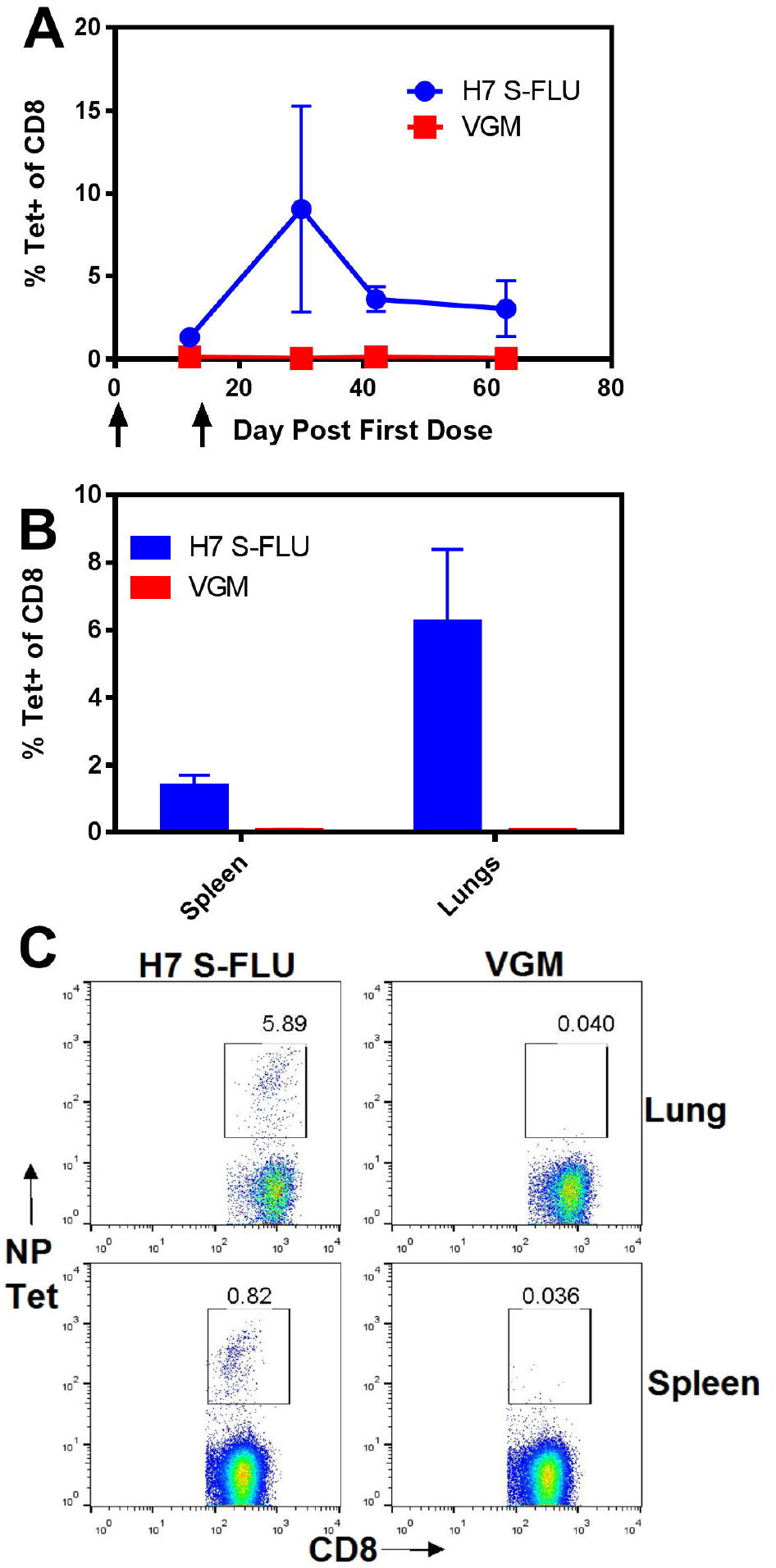
H7 S-FLU Intranasal immunization induces CD8+ T cells in lungs, spleen and blood. (A) C57BL/6 mice were immunized with 16 HAU H7 (7.35×10^5^ TCID_50_) S-FLU or VGM at days 0 and 15 and blood (n=4-10) was collected at days 12, 30, 42 and 63: NP_366–74_ specific CD8+ T cells were quantified by tetramer staining. Values are means +/-SD. (B) On day 64 (n=4) mice were killed, organs were collected and the percentage of NP_366–74_ specific CD8+ T cells determined by FACS. Data are mean +/-SD. (C) FACS of percent tetramer positive cells within a CD8 gated population on day 64 from one representative animal. Data are from one representative experiment with 10 mice per group.

#### Antibody to HA

We have previously shown that after lung vaccination with doses of < 10^6^ TCID_50_ a weak antibody response to the HA pseudotyping (coating) the S-FLU particle is generated that can be detected by ELISA but rarely in the MN assay [43, 46]. This was the case at day 64, 40 days after the second dose of 16 HAU (7×10^5^ TCID_50_), H7-S-FLU neutralising antibody was not detectable in serum (Figure 3, middle panel, column 7). Mice given 32 HAU (5×10^6^ TCID_50_) of an H5 S-FLU, produced a weak MN response to H5 (Figure 3, middle panel, column 8). By contrast i.p. priming with a ten-fold greater dose generated high levels of specific neutralising antibody to both H7 HA and the encoded N1 NA (Figures 3–5)

**Figure 3:**
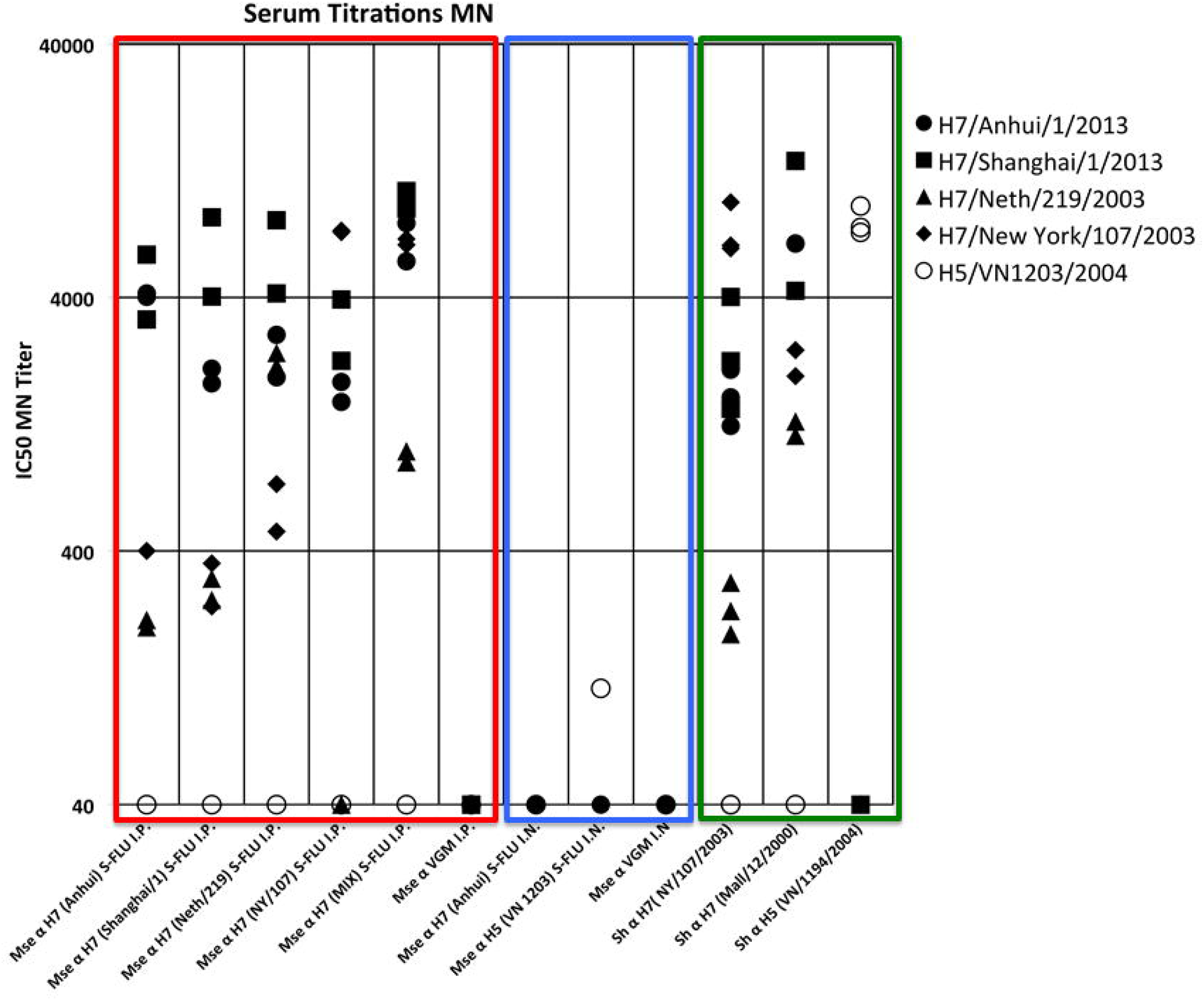
Intraperitoneal, but not intranasal vaccination with S-FLU induces neutralising antibody to the H7 HA coating the S-FLU particle. Columns 1-6 (boxed in red): C57BL/6 male mice were immunized twice i.p. with various H7 S-FLUs on days 0 and 28 and bled on day 42. Sera from 6 mice were pooled. The H7 S-FLU doses were 160 HAU Anhui/1/2013, 40 HAU Shanghai/1/2013, 1280 HAU Netherlands/219/203, 640 HAU New York/107/2003 S-FLU or a Mixture of all four. Columns 7-9 (boxed in blue): in a separate experiment (shown in Figure 2) female C57BL/6 mice were immunized on days 0 and 15, in the lung, with S-FLU (16 HAU H7 Anhui/1/2013 or 32 HAU H5 VN1203) and bled on day 64. Sera from 4 mice were pooled. Columns 10-13 (boxed in green): Control antibodies consisting of sheep antisera to H7 proteins: A/New York/107/2003 (NIBSC 09/148), and to A/mallard/Netherlands/12/2000 (NIBSC 07/278); sheep antiserum to H5 A/VN/1194/2004 (NIBSC 07/148). Each IC_50_ point is derived from a titration curve done in duplicate as described in methods.

After two doses of 160 HAU = 7×10^6^ TCID_50_ H7 (Anhui/1/2013) S-FLU virus i.p. titres of neutralising HA specific antibody were induced (Figure 3, column 1) equivalent in titre to the hyperimmune standard control sheep sera. To assess whether responses to PR8 internal genes or their products contributed to the neutralising titre, and to study specificity, we tested neutralisation of an H5 S-FLU virus that differs only in the HA coating the particle. The serum raised against H7 S-FLU showed no neutralisation on H5 S-FLU in the MN assay (Figure 3, column 1). We further characterized the antibody generated by Anhui H7 S-FLU for the degree of cross neutralisation of S-FLU coated with related H7 HAs from viruses known to have caused human infections. Pooled sera raised against two i.p. doses neutralised the Anhui H7 S-FLU with an IC_50_ titre of ~1:4000 and showed a similar titre to the co-circulating strain A/Shanghai/1/2013, but reduced titres (≤ 400) to A/New York/107/2003 and A/Netherlands/219/2003.

In parallel we induced antibodies by i.p. immunization twice with each of the four H7 S-FLU strains individually, and against a mixture of all four. Due to differences in yield, various doses ranging between 40-1280 HAU were used. Each strain of H7 S-FLU generated strong neutralising antibody against itself (Figure 3, columns 1-4). Notably, two 40 HAU doses of A/Shanghai/1/2013 H7 S-FLU induced a titre nearly as high as the other three, which were given at higher doses of 160-1280 HAU. The individual antisera showed varying cross-reactivity with H7 HAs from more distantly related viruses, with the antiserum raised to 1280 HAU of A/Netherlands/219/2003 HA showing the broadest reactivity. Antiserum raised to S-FLU coated in H7 A/New York/107/2003 (North American origin) failed to neutralise A/Netherlands/219/2003 (Eurasian origin), but did cross-react with A/Anhui/1/2013 and A/Shanghai/1/2013. The polyvalent vaccine mix of four H7 S-FLUs induced strong neutralising titres to all four of the immunizing viruses (Figure 3, column 5). Mice immunized with VGM (no virus) did not produce H7-neutralising antibody (Figure 3, column 6).

A control hyperimmune sheep antiserum raised to A/New York/107/2003 HA (NIBSC 09/148) showed very strong neutralisation of the A/New York/107/2003 H7 S-FLU, moderate cross-reactivity with the A/Anhui/1/2013 and A/Shanghai/1/2013 H7 S-FLU viruses, and weak activity with A/Netherlands/219/2003 (Figure 3, column 10). A second control sheep serum raised to A/mallard/Netherlands/12/2000 (NIBSC 07/278) neutralised all four H7 S-FLU strains (Column 11). Neither serum raised to H7 viruses neutralised the H5 S-FLU and the H5 specific sheep antiserum strongly neutralised the H5 S-FLU but had no effect on any of the H7 S-FLU viruses (Figure 3, columns 10-12).

The above results were obtained in C57BL/6 mice. In addition, BALB/c mice were immunized twice i.p. with H7 S-FLU Anhui/1/2013 with resultant neutralising antibody to H7 S-FLU (Figure 4A) and a strong CTL response in spleen (Figure 4B). Neutralisation of four H7 S-FLU strains by the BALB/c antiserum to A/Anhui/1/2013 was tested (Figure 4C), and results were similar to C57BL/6 (Figure 3) but the BALB/c antiserum to A/Anhui/1/2013 showed no cross-reactivity with A/Netherlands/219/2003.

**Figure 4:**
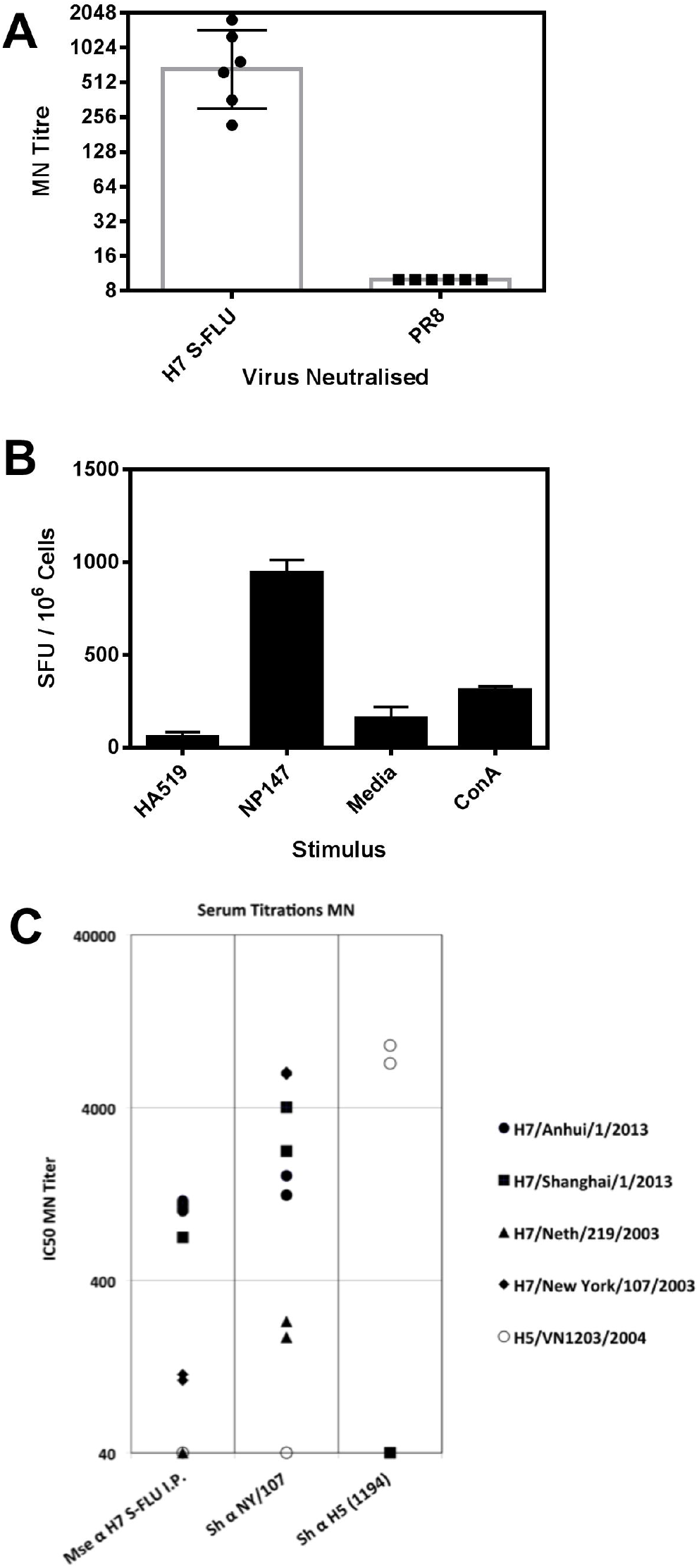
H7 S-FLU induces neutralising antibody and splenic T cells in vivo after i.p. immunization of BALB/c mice. Animals were primed with 160 HAU H7 (A/Anhui/1/2013) S-FLU or VGM on day 0 and day 14 and samples were collected at day 66. (A) MN titres to H7 S-FLU (Anhui/1/2013) and PR8 were determined for individual mice. Values are geometric mean +/-geometric SD. (B) ELISPOT assay of CD8 T cells from spleen showing responses to conserved peptide NP147 and controls at day 66 post i.p. priming. Data are mean +/-SD. (C) Pooled antisera from the 6 mice represented in (A) were tested for cross neutralisation of S-FLU pseudotyped with various H7 HAs derived from A/Anhui/1/2013, A/Shanghai/1/2013, A/Netherlands/219/2003, A/New York/107/2003, with H5 A/VietNam/1203/2004 S-FLU as a specificity control. Control antisera were from a sheep hyper-immunised with H7 A/New York/107/2003 (NIBSC 09/148) or H5 A/VietNam/1194/2004 (NIBSC 07/148). Each IC_50_ point is derived from a titration curve done in duplicate.

#### Antibody to NA

In contrast to HA, S-FLU encodes a viable NA vRNA and when cells are infected with S-FLU NA protein is expressed at the cell surface at a similar level to PR8 wt virus (Figure 1). We measured the neutralising response to the encoded PR8 NA by ELLA, using either PR8 wt virus or an H5 S-FLU as sources of N1 NA activity in the assay to avoid possible interference by antibodies to H7 HA [60, 61]. To replicate the results we immunized mice with an H5 S-FLU in parallel and measured the NA response on PR8 wt virus as a source of matched NA. Figure 5 (columns 7-9) shows that lung vaccination with either H7 (Anhui/1/2013) or H5 (VN/1203/2004) S-FLU induced a significant titre (IC_50_ > 1:1000) of neutralising antibody to N1 (PR8) NA that was strain specific and did not cross-neutralise the N1 NA from A/Eng/195/2009 or N2 from X31 (A/Aichi/2/1968). Intraperitoneal vaccination with a ten-fold higher dose of S-FLU induced a higher titre to ~1:4:000 (Figure 5 columns 2-5 compared to columns 7–8).

**Figure 5.**
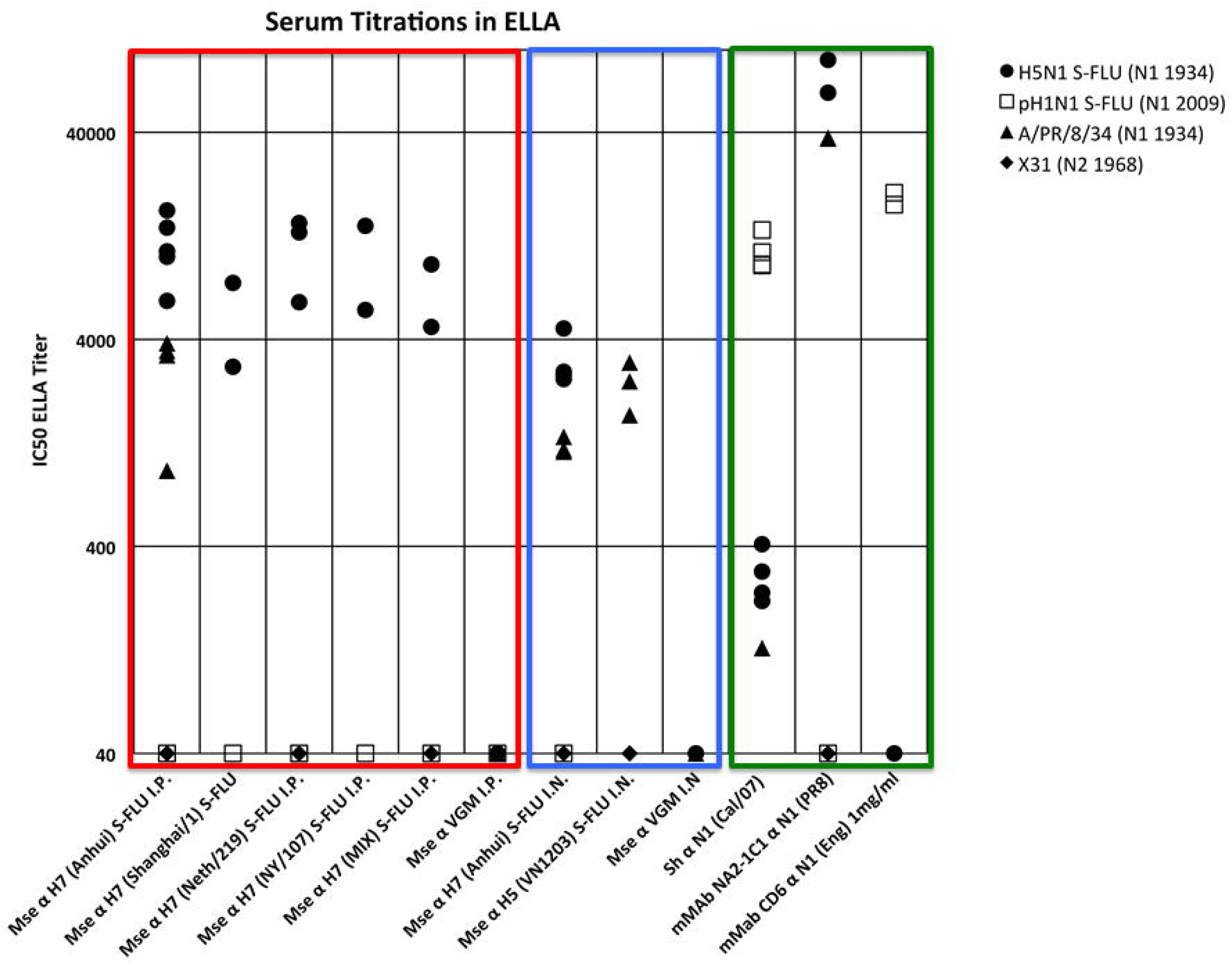
Intraperitoneal and Intra-lung vaccination with S-FLU induces neutralising antibody to the N1 NA. Columns 1-9: as for Figure 3. Columns 10-12: control antibodies consisting of sheep antiserum to N1 from A/California/07/2009 (NIBSC 10/218), murine mAb NA2-1C1 specific for N1 from A/PR/8/34, murine mAb CD6 specific for N1 from A(H1N1)pdm09 influenza. Each IC_50_ point is derived from a titration curve done in duplicate.

Control antibodies included a sheep antiserum to N1 from A/California/07/2009 (NIBSC 10/218) which showed a high titre to the closely related A/Eng/195/2009 N1pdm09 NA and a weak cross-reaction with PR8 N1 NA (Figure 5, column 10).

Additional specificity controls were (i) the murine mAb NC2-1C1 [53] that inhibited the N1 from PR8 but not N1pdm09 (Column 11), ii) the murine mAb CD6 [54] that inhibited viruses carrying N1pdm09 NA with an IC_50_ of 49 ng/mL, similar to the reported value of 64 ng/mL [54], but was non-reactive with the N1 (PR8), (Figure 5, column 12).

### Cross protection following vaccination with H7 S-FLU

To assess if H7 S-FLU vaccination gave cross protection, groups of mice were immunized in the lung with H7 S-FLU virus twice at two weekly intervals and then challenged one month later with 0.32 HAU (10^4^ TCID_50_) of PR8 virus. Immunized mice were protected against PR8 challenge (Figure 6A) whilst VGM treated mice succumbed with rapid weight loss and clinical signs of morbidity (Figure 6A, B & F). Similarly, mice vaccinated with 16 HAU of H7 S-FLU were also protected against 32 HAU of X31 (H3N2) virus challenge (Figure 6B), to which there was no neutralising antibody to H3 or N2 (Data not shown). Neither immunized nor control mice were protected against influenza B virus challenge (Figure 6C, F).

**Figure 6.**
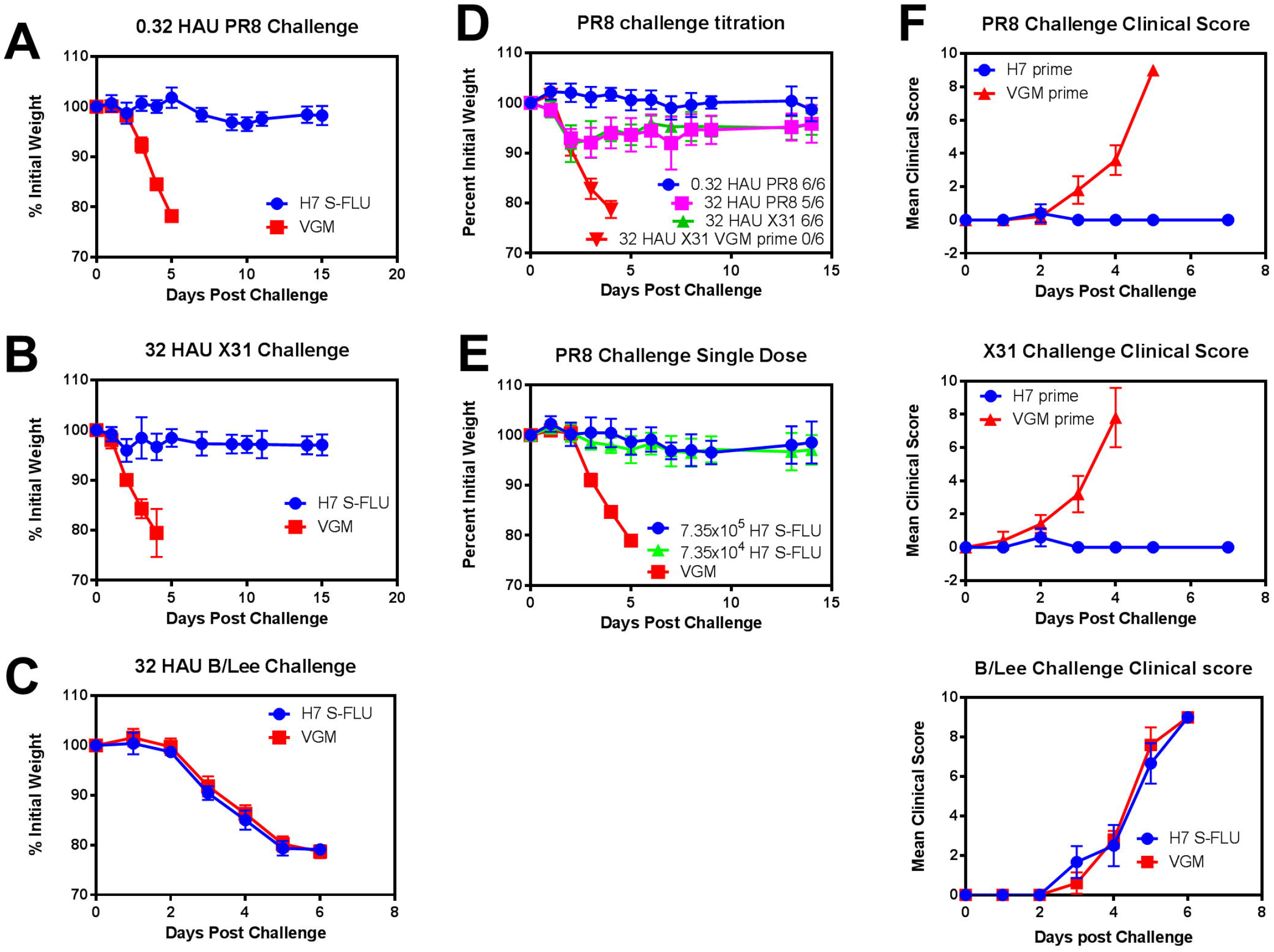
H7 S-FLU vaccination protects against challenge with X31 (H3N2) and PR8 (H1N1) viruses but not influenza B virus. BALB/c mice were primed with 16 HAU (7.35×10^5^ TCID_50_) H7 S-FLU or VGM on day 0 and day 14 (A-D) or day 14 only with 7.35×10^5^ or 7.35×10^4^ TCID_50_ (E) and then challenged with (A) 0.32 HAU (10^4^ TCID_50_) PR8, (B) 32 HAU X31, (C) 32 HAU B/Lee virus, (D) up to 32 HAU (10^6^ TCID_50_) PR8 or (E) 0.32 HAU PR8 virus at least one month later and monitored daily for weight loss. (F) BALB/c mice primed with H7 S-FLU or VGM at days 0 and 14 and challenged with 0.32 HAU PR8 (top), 32 HAU X31 (middle) or 32 HAU B/Lee (bottom) on day 39 and monitored for clinical score. All data are from one representative experiment repeated at least twice. Data shown are mean +/-SD.

When a challenge dose of 0.32 HAU (10^4^ TCID_50_) PR8 was used in H1 (PR8) S-FLU primed animals, all (6/6) were protected while 5/6 animals were protected against the full dose of 32 HAU (~10^6^ TCID_50_ = >10^4^ LD50) PR8, but with more weight loss than the 0.32 HAU PR8 challenge (Figure 6D). This level of protection was less than that seen with homotypic challenge after H1 (PR8) S-FLU priming and wild type PR8 challenge, where complete protection without weight loss was observed [43]. In addition a single dose of 7.35×10^5^ or 7.35×10^4^ TCID_50_ H7 S-FLU was able to protect mice against challenge with 0.32 HAU (10^4^ TCID_50_) PR8 (Figure 6E).

### Titration of H7 S-FLU and histology during challenge

Further to showing that a dose of 16 HAU (7.35×10^5^ TCID_50_) of H7 S-FLU was protective, we determined at which concentration two doses of the H7 S-FLU vaccine failed to protect. Animals were dosed with ten-fold dilutions of H7 S-FLU and doses as low as ~700 TCID_50_ were protective while a dose of ~70 TCID_50_ was not (Figure 7A). However, mice immunized twice with 735 TCID_50_ did have more noticeable weight loss upon challenge than mice immunized with higher doses (Figure 7A). We also measured virus titre in the lungs at day 3 post-challenge for some of these groups: mice primed with two doses of 7×10^5^ and 7×10^4^ TCID_50_ of H7 S-FLU efficiently cleared virus by day 3 while 7 TCID_50_ and VGM primed mice had high virus titres in lungs (Figure 7B). These data, and a review of the literature on protection by LAIV, would suggest that a reliable priming dose to the lung for heterotypic protection in mice is 10^4^ – 10^6^ TCID_50_ of S-FLU.

**Figure 7.**
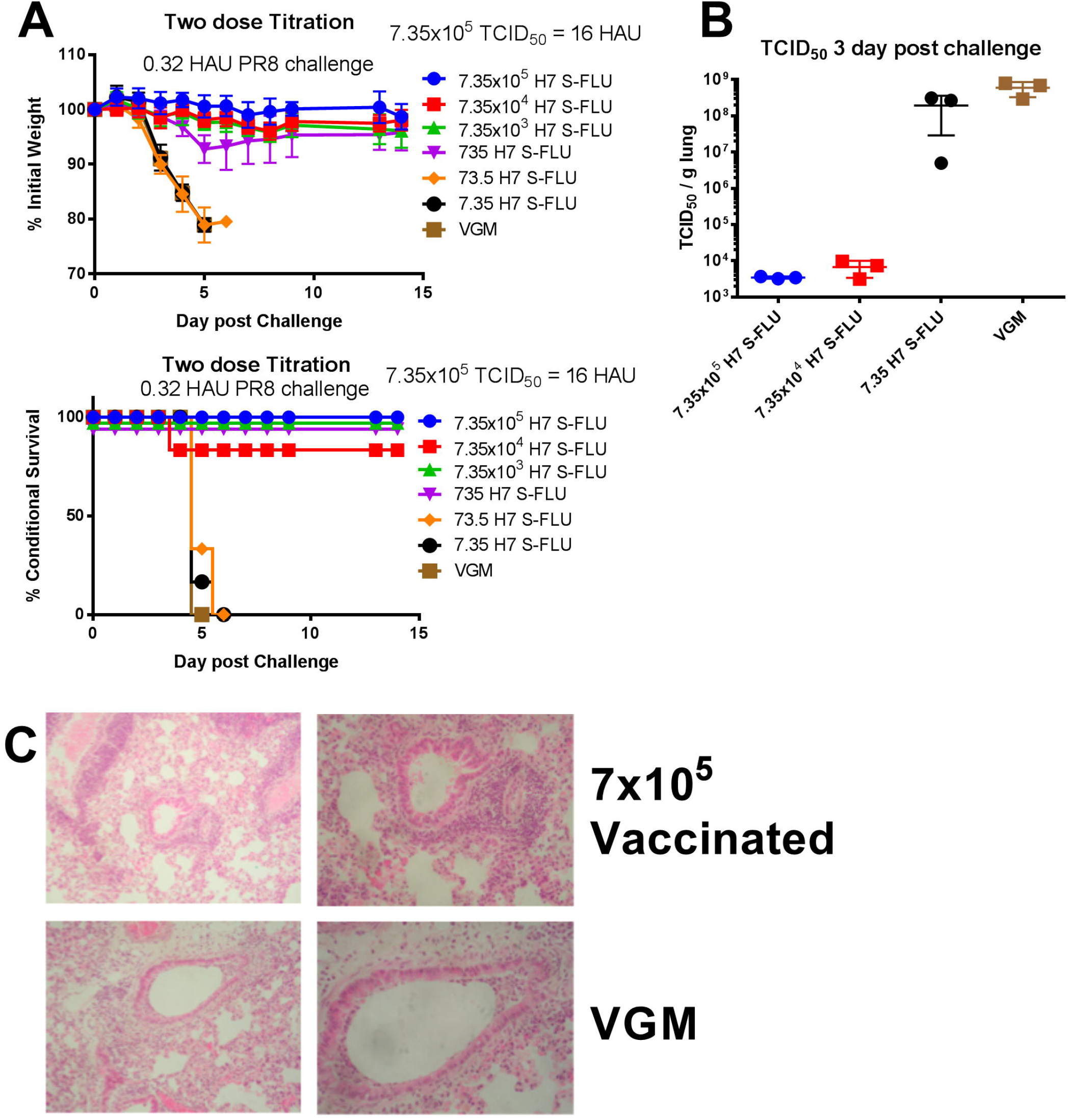
Titration of H7 S-FLU priming dose reveals a protection threshold of 735 TCID_50_ of H7 S-FLU. BALB/c mice were primed with VGM or titrated doses of H7 S-FLU at day 0 and day 15 and challenged with 0.32 HAU [10^4^ TCID_50_] PR8 virus on day 49. (A) Weight loss and survival were measured. Mean +/-SD shown. (B) Lungs were collected from killed mice at day 3 post-challenge and virus titre measured with mean and SD shown; (C) tissue sections cut and stained with hematoxylin/eosin (Left panel x25, Right x40) from mice primed twice with 7×10^5^ TCID_50_ H7 S-FLU or VGM (control mice). Note i) the perivascular cuffs of infiltrating lymphocytes in the bronchovascular bundle and preservation of the columnar epithelium in the small bronchioles in the sample from the vaccinated animal and ii) necrotic debris in the lumen of the small bronchiole and destruction of columnar epithelium in the sample from a control mouse. Data are from one representative experiment with at least 6 mice per group

The histology of the lungs of mice primed twice with 7×10^5^ TCID_50_ H7 S-FLU or VGM primed mice, 3 days after challenge, was compared. H7 S-FLU immunized mice at day 3 post challenge showed aggregation of lymphocytes surrounding the venules in the bronchovascular bundle that were not present in control virus challenged mice, with conservation of the columnar epithelium in the small bronchioles. By contrast the sections from control animals did not contain a perivascular cuff of lymphocytes and showed destruction of the columnar epithelium with accumulation of necrotic debris in the small bronchioles (necrotizing bronchiolitis) (Figure 7C)

### Post challenge titres of antibody

To establish whether neutralising antibody and T cells were generated to the priming and challenge doses given i.n, we performed MN and ELISA assays on serum samples from mice collected at least 30 days post-challenge. A significant response to H7 HA was detected by ELISA (not shown), but the antibodies were not neutralising (Figure 3), which is consistent with responses to the H1 S-FLU described previously [43]. By contrast there was a strong response to the challenge virus in the serum (Figure 8).

**Figure 8.**
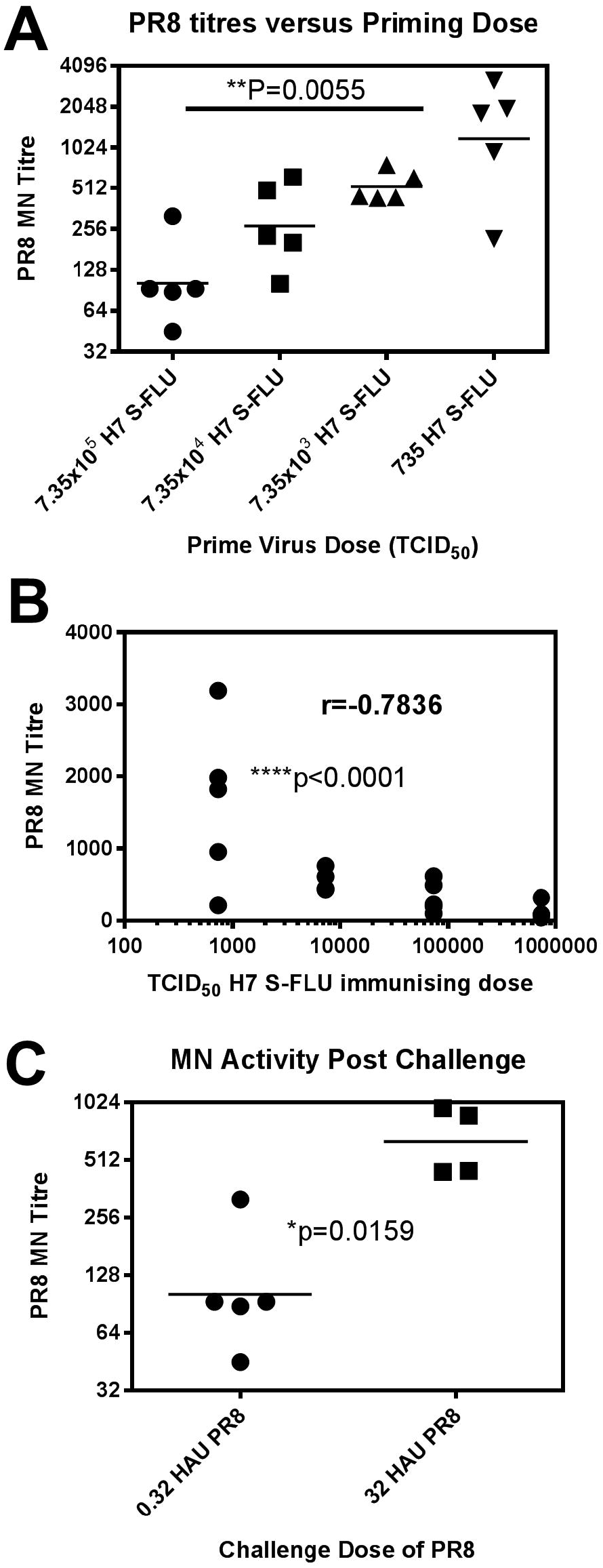
The titre of neutralising antibody generated against the challenge influenza virus is inversely related to the vaccination dose of H7 S-FLU, and directly related to the PR8 challenge dose. BALB/c mice were primed in the lung with VGM or titrated doses of H7 S-FLU at day 0 and day 15, challenged on day 49 with 0.32 HAU of PR8 virus, and allowed to recover from challenge. (A) Serum MN antibody titre was measured on day 97 samples. One way ANOVA with multiple correction p value comparing 7.35×10^5^ and 735 TCID_50_ groups shown and (B) correlation of MN titre with priming dose of H7 S-FLU. Spearman correlation r value and p value shown. (C) Comparison of MN titres in serum at day 97 between mice challenged with 0.32 or 32 HAU PR8 after a vaccination dose of 7×10^5^ TCID_50_ of H7 S-FLU. These data are from one experiment repeated twice. Means shown are geometric means.

The neutralising anti-HA antibody response to PR8 challenge virus was dependent on both the priming dose of H7 S-FLU and the challenge dose of PR8. Figure 8A shows that the MN titre in the serum to the PR8 challenge virus was inversely proportional to the priming dose of H7 S-FLU (Figure 8B). In addition, mice challenged with 32 HAU of PR8 had higher MN titres to PR8 than mice challenged with 0.32 HAU PR8 (Figure 8C).

## Discussion

We have shown that a new version of S-FLU coated in H7 HA from A/Anhui/1/2013 can generate heterotypic protective immune responses to A/PR/8/34 (H1N1) and X31 (H3N2) when administered to the mouse lung as a single dose of 7×10^4^ TCID_50_ or two doses between 7×10^2^ – 7×10^5^ TCID_50_. The protective effect is associated with a strong class I restricted T cell response to conserved peptides derived from the virus NP, a moderate subtype specific antibody response to NA, but a weak neutralising antibody response to the H7 HA used to pseudotype (coat) the particles. H7 (Anhui/1/2013)-S-FLU propagated to a titre of 1–2 × 10^7^ TCID_50_/ml, and H7 (Netherlands/219/2003)-S-FLU gave >10 fold greater yields up to 3 × 10^8^ TCID_50_/mL, similar to the A/PR/8/34 parental virus. The A/Netherlands/219/2003 wt virus replicated to higher levels in murine tissues and induced the broadest antibody response of several H7 viruses [65], as also shown in Figure 3 for the equivalent S-FLU. If S-FLU were to be given to humans in a similar dosage to LAIV this represents 1–10 doses/ml for human and 1–10 × 10^4^ murine doses/mL. These results confirm and extend our earlier experiments with S-FLU coated in H1 HAs [43], and the work on related single-cycle infectious influenza A viruses (reviewed in [46]).

The protective effect of the H7 S-FLU is specific to influenza A (not B), as found in the classic description of “partial specific heterotypic immunity” [11], which was transferrable with cross-reactive class I restricted CTL [24–26]. Evidence has accumulated for a cross protective effect associated with the Class I restricted T cell response in humans [35, 36], including recent evidence for a correlation between a strong cross-reactive CTL response and milder outcome after H7N9 infection in humans [39]. We also showed that an H5 S-FLU with internal genes from A/PR/8/34 can prevent replication of wild type H7N9 (A/Anhui/1/2013) in mice and ferrets [44]. This suggests that S-FLU and other single-cycle infectious influenza A viruses [46] could be useful in the stimulation of cross subtype immunity for protection against future pandemic influenza A viruses.

Antibodies to NA can have a significant protective effect against influenza in animal models of infection [66, 67], and evidence from vaccine trials strongly suggests this is true for human infections [68]. The inhibition of NA by antibody does not usually block virus entry (so would not be detected in the neutralisation assay we use), but prevents virus release, and reduces plaque size in vitro [54, 66, 69]. Recent evidence shows that an engineered influenza can use NA as a receptor-binder, making the sialic acid binding function of HA redundant, and reserving HA for its fusion function alone [70, 71]. If such a virus evolved naturally, vaccines that stimulate a strong neutralising antibody to NA would be essential.

We have demonstrated that cells infected with S-FLU express the PR8 NA in the absence of HA at the cell surface at similar levels to cells infected with wt A/PR/8/34 (Figure 1). Therefore S-FLU might be expected to induce an antibody response to NA, and we have shown this conclusively, both after administration to the lung at low dose, and an enhanced response after larger doses i.p. (Figure 5). The response to NA is strain specific, as the antibody induced to the N1 NA from PR8 does not cross-inhibit the N1 NA from A/Eng/195/2009 or the N2 from X31 (A/Aichi/2/68), and this was the case also for the control mAb to N1 (PR8), NA2-1C1. It is likely that the anti N1 antibody response may have contributed to protection from A/PR/8/34, but is unlikely to have contributed to protection from the fully mismatched X31 (H3N2). However cross-reactivity between the recent H1N1pdm09 N1 neuraminidase and the H5N1 strains [72–74] could be harnessed to add a useful additional layer of protection induced by S-FLU where relevant. Future work will reveal if it is feasible to immunize with a mix of NAs in the S-FLU to induce a broadly reactive NA response.

After i.n. administration of a low dose (<10^6^ TCID_50_) of S-FLU an antibody response to the HA coating S-FLU can be detected by ELISA [43], but it is predominantly non-neutralising (Figure 3). This is probably due to the lack of synthesis of new HA by S-FLU infected cells. However, H7 S-FLU given at ten-fold higher dose, in the peritoneal space, generates a robust neutralising antibody titre to A/Anhui/1/2013 HA, in addition to a T cell response to a conserved nucleoprotein epitope detected in the spleen (Figures 3, 4). The dose required to induce this neutralising antibody response, while ten-fold higher than that given to the lung, is still a low dose in absolute terms: 160 HAU (~7 ×10^6^ TCID_50_) is expected to contain less than 10 ng of HA glycoprotein. Three additional strains of H7 S-FLU induced strong neutralising responses to themselves, including a fourfold lower dose (40 HAU) of the A/Shanghai/1/2013 H7 S-FLU, which did not grow to as high a titre as the others.

The antibody induced to H7 HA was relatively strain specific. However peripheral vaccination with a mix of the four strains of H7 S-FLU, which included North American and Eurasian strains, induced a strong neutralising response to all the viruses in the mix. Peripheral immunization with a polyvalent S-FLU, that can induce neutralising antibodies to multiple strains, might be an alternative means of inducing broadly neutralising antibody to HA, as opposed to efforts to induce antibodies to the conserved stem region of the HA. This principle has been shown for a trivalent H5 DNA vaccine [75], and a tetravalent VLP vaccine [76]. These new results with an H7 from a group 2 virus, that is viewed as a pandemic threat, extend our earlier work with S-FLU coated in H1 HAs from group 1 viruses, A/PR/8/34 and the H1N1pdm09 virus A/England/195/2009, which also induced strain specific neutralising antibodies after i.p. inoculation [43]. This suggests that S-FLU particles are strongly immunogenic for B cells when given in the periphery, as has been seen with wt influenza [77].

In vaccinated mice we found a strong neutralising antibody response to the challenge virus at one month post infection, in situations where replication of the challenge virus was clearly suppressed at day 3 post-challenge (Figures 7,8). The titre of neutralising antibody generated post-challenge was inversely proportional to the dose of the priming vaccine S-FLU virus vaccine, as was the degree of replication of the challenge virus detected in the lung. This suggests that the level of neutralising antibody generated after challenge is related to the degree of replication that occurs, and the higher dose of H7 S-FLU gives a stronger immune response that prevents challenge virus replication and thus dampens the antibody response. The animals which received lower doses of vaccine had detectable virus replication at day 3, lost more weight, and generated the highest neutralising antibody titres to the challenge virus.

The heterotypic protection provided by S-FLU is thus not sterilizing, as indicated in the first description of this phenomenon [11]. The strong neutralising response to the challenge virus may be an additional benefit of S-FLU, as this would provide longterm antibody-based protection, and also strengthen and prolong the T cell response.

Together these results show that S-FLU is a flexible influenza vaccine candidate as it can be produced with various surface HAs and NAs, it can stimulate both local cross protective T cell responses, and systemic neutralising antibodies to HA and NA, depending on how it is administered. In principle, it could be given both to the lung to induce cross protective T cells, and as a mix of strains in the periphery to induce both a broad neutralising antibody response and sustain the heterosubtypic protection provided by the T cell response [21].

## Acknowledgements

We wish to thank John Paul Jukes, Dawn Shepherd and Daniel Puleston for synthesis and quality control of the NP366 tetramers. We also thank staff at the BMS for care and maintenance of animal colonies. We are grateful for the provision of tissue sections by the Oxford Centre for Histocompatibility Research OUH NHS Trust Oxford. We thank Maryna Eichelberger and Jon Yewdell for antibodies. These studies were supported by the Townsend-Jeantet Charitable Trust (registered charity 1011770), Oxford University and the MRC Human Immunology Unit.

## Author Contributions

TJP and ARMT: designed and undertook the experiments, collected and analysed data, structured, wrote and edited paper. HB, RMS, PR, MH, K-YH, RD & LS performed experiments, collected and analysed data.

## Competing Interests Statement

ARMT is named on a patent for S-FLU as a vaccine, ownership of which is shared between Oxford University and The Townsend-Jeantet Charitable Trust (registered charity 1011770). All other authors have no competing interests to declare.

## References

1. Medina, R.A. and A. Garcia-Sastre, Influenza A viruses: new research developments. Nature Reviews Microbiology, 2011. 9(8): p. 590–603.

2. Fouchier, R.A., et al., Avian influenza A virus (H7N7) associated with human conjunctivitis and a fatal case of acute respiratory distress syndrome. Proc Natl Acad Sci U S A, 2004. 101(5): p. 1356–61.

3. Peiris, M., et al., Human infection with influenza H9N2. Lancet, 1999. 354(9182): p. 916–7.

4. Gao, R., et al., Human infection with a novel avian-origin influenza A (H7N9) virus. N Engl J Med, 2013. 368(20): p. 1888–97.

5. Xiong, X., et al., Receptor binding by an H7N9 influenza virus from humans. Nature, 2013. 499(7459): p. 496–9.

6. Hancock, K., et al., Cross-reactive antibody responses to the 2009 pandemic H1N1 influenza virus. N Engl J Med, 2009. 361(20): p. 1945–52.

7. Hardelid, P., et al., Assessment of baseline age-specific antibody prevalence and incidence of infection to novel influenza A/H1N1 2009. Health Technol Assess, 2010. 14(55): p. 115–92.

8. Huang, K.Y., et al., Focused antibody response to influenza linked to antigenic drift. J Clin Invest, 2015. 125(7): p. 2631–45.

9. Paules, C.I., et al., The Pathway to a Universal Influenza Vaccine. Immunity, 2017. 47(4): p. 599–603.

10. Smith, W., C.H. Andrewes, and P.P. Liadlaw, A virus obtained from influenza patients. Lancet, 1933. 2: p. 66–8.

11. Schulman, J.L. and E.D. Kilbourne, Induction of Partial Specific Heterotypic Immunity in Mice by a Single Infection with Influenza a Virus. J Bacteriol, 1965. 89: p. 170–4.

12. Doherty, P.C. and A. Kelso, Toward a broadly protective influenza vaccine. J Clin Invest, 2008. 118(10): p. 3273–5.

13. Epstein, S.L. and G.E. Price, Cross-protective immunity to influenza A viruses. Expert Rev Vaccines, 2010. 9(11): p. 1325–41.

14. Schulman, J.L., Experimental transmission of influenza virus infection in mice. 3. Differing effects of immunity induced by infection and by inactivated influenza virus vaccine on transmission of infection. J Exp Med, 1967. 125(3): p. 467–78.

15. Steel, J., et al., Transmission of pandemic H1N1 influenza virus and impact of prior exposure to seasonal strains or interferon treatment. J Virol, 2010. 84(1): p. 21–6.

16. Fazekas de St Groth, S. and M. Donnelly, Studies in experimental immunology of influenza. IV. The protective value of active immunization. Aust. J. Exp. Biol. Med. Sci., 1950. 28: p. 61–75.

17. Nguyen, H.H., et al., Heterosubtypic immunity to lethal influenza A virus infection is associated with virus-specific CD8(+) cytotoxic T lymphocyte responses induced in mucosa-associated tissues. Virology, 1999. 254(1): p. 50–60.

18. Perrone, L.A., et al., Intranasal vaccination with 1918 influenza virus-like particles protects mice and ferrets from lethal 1918 and H5N1 influenza virus challenge. J Virol, 2009. 83(11): p. 5726–34.

19. Lau, Y.F., A.R. Wright, and K. Subbarao, The contribution of systemic and pulmonary immune effectors to vaccine-induced protection from H5N1 influenza virus infection. J Virol, 2012. 86(9): p. 5089–98.

20. Wu, T., et al., Lung-resident memory CD8 T cells (TRM) are indispensable for optimal cross-protection against pulmonary virus infection. J Leukoc Biol, 2014. 95(2): p. 215–24.

21. Uddback, I.E., et al., Combined local and systemic immunization is essential for durable T-cell mediated heterosubtypic immunity against influenza A virus. Sci Rep, 2016. 6: p. 20137.

22. Zweerink, H.J., et al., Cytotoxic T cells kill influenza virus infected cells but do not distinguish between serologically distinct type A viruses. Nature, 1977. 267(5609): p. 354–6.

23. Effros, R.B., et al., Generation of both cross-reactive and virus-specific T-cell populations after immunization with serologically distinct influenza A viruses. J Exp Med, 1977. 145(3): p. 557–68.

24. Yap, K.L., G.L. Ada, and I.F. McKenzie, Transfer of specific cytotoxic T lymphocytes protects mice inoculated with influenza virus. Nature, 1978. 273(5659): p. 238–9.

25. Lin, Y.L. and B.A. Askonas, Biological properties of an influenza A virus-specific killer T cell clone. Inhibition of virus replication in vivo and induction of delayed-type hypersensitivity reactions. J Exp Med, 1981. 154(2): p. 225–34.

26. Taylor, P.M. and B.A. Askonas, Influenza nucleoprotein-specific cytotoxic T-cell clones are protective in vivo. Immunology, 1986. 58(3): p. 417–20.

27. Townsend, A.R. and J.J. Skehel, The influenza A virus nucleoprotein gene controls the induction of both subtype specific and cross-reactive cytotoxic T cells. J Exp Med, 1984. 160(2): p. 552–63.

28. Townsend, A.R., F.M. Gotch, and J. Davey, Cytotoxic T cells recognize fragments of the influenza nucleoprotein. Cell, 1985. 42(2): p. 457–67.

29. Yewdell, J.W., et al., Influenza A virus nucleoprotein is a major target antigen for cross-reactive anti-influenza A virus cytotoxic T lymphocytes. Proc Natl Acad Sci U S A, 1985. 82(6): p. 1785–9.

30. Townsend, A.R., et al., The epitopes of influenza nucleoprotein recognized by cytotoxic T lymphocytes can be defined with short synthetic peptides. Cell, 1986. 44(6): p. 959–68.

31. Liang, S., et al., Heterosubtypic immunity to influenza type A virus in mice. Effector mechanisms and their longevity. J Immunol, 1994. 152(4): p. 1653–61.

32. Epstein, S.L., et al., Mechanisms of heterosubtypic immunity to lethal influenza A virus infection in fully immunocompetent, T cell-depleted, beta2-microglobulin-deficient, and J chain-deficient mice. J Immunol, 1997. 158(3): p. 1222–30.

33. Laidlaw, B.J., et al., Cooperativity between CD8+ T cells, non-neutralizing antibodies, and alveolar macrophages is important for heterosubtypic influenza virus immunity. PLoS Pathog, 2013. 9(3): p. e1003207.

34. McMichael, A.J., et al., Declining T-cell immunity to influenza, 1977-82. Lancet, 1983. 2(8353): p. 762–4.

35. Sridhar, S., et al., Cellular immune correlates of protection against symptomatic pandemic influenza. Nature medicine, 2013. 19(10): p. 1305–12.

36. Hayward, A.C., et al., Natural T Cell Mediated Protection Against Seasonal and Pandemic Influenza: Results of the Flu Watch Cohort Study. Am J Respir Crit Care Med, 2015. 191(12): p. 1422–31.

37. van de Sandt, C.E., et al., Human cytotoxic T lymphocytes directed to seasonal influenza A viruses cross-react with the newly emerging H7N9 virus. J Virol, 2014. 88(3): p. 1684–93.

38. Lee, L.Y., et al., Memory T cells established by seasonal human influenza A infection cross-react with avian influenza A (H5N1) in healthy individuals. J Clin Invest, 2008. 118(10): p. 3478–3490.

39. Wang, Z., et al., Recovery from severe H7N9 disease is associated with diverse response mechanisms dominated by CD8(+) T cells. Nat Commun, 2015. 6: p. 6833.

40. McMichael, A.J., et al., Cytotoxic T-cell immunity to influenza. N Engl J Med, 1983. 309(1): p. 13–7.

41. Wilkinson, T.M., et al., Preexisting influenza-specific CD4+ T cells correlate with disease protection against influenza challenge in humans. Nat Med, 2012. 18(2): p. 274–80.

42. Townsend, A.R., et al., Cytotoxic T lymphocytes recognize influenza haemagglutinin that lacks a signal sequence. Nature, 1986. 324(6097): p. 575–7.

43. Powell, T.J., et al., Pseudotyped influenza A virus as a vaccine for the induction of heterotypic immunity. J Virol, 2012. 86(24): p. 13397–406.

44. Baz, M., et al., Nonreplicating influenza A virus vaccines confer broad protection against lethal challenge. MBio, 2015. 6(5): p. e01487–15.

45. Morgan, S.B., et al., Aerosol Delivery of a Candidate Universal Influenza Vaccine Reduces Viral Load in Pigs Challenged with Pandemic H1N1 Virus. J Immunol, 2016. 196(12): p. 5014–23.

46. Nogales, A., et al., Development and applications of single-cycle infectious Influenza A Virus (sciIAV). Virus Res, 2015.

47. Chen, Z., et al., Development of a high-yield live attenuated H7N9 influenza virus vaccine that provides protection against homologous and heterologous H7 wild-type viruses in ferrets. J Virol, 2014. 88(12): p. 7016–23.

48. Jin, H. and K. Subbarao, Live attenuated influenza vaccine. Curr Top Microbiol Immunol, 2015. 386: p. 181–204.

49. Mallory, R.M., T. Yi, and C.S. Ambrose, Shedding of Ann Arbor strain live attenuated influenza vaccine virus in children 6-59 months of age. Vaccine, 2011. 29(26): p. 4322–7.

50. Matrosovich, M., et al., Overexpression of the alpha-2,6-sialyltransferase in MDCK cells increases influenza virus sensitivity to neuraminidase inhibitors. J Virol, 2003. 77(15): p. 8418–25.

51. Kilbourne, E.D., Future influenza vaccines and the use of genetic recombinants. Bull World Health Organ, 1969. 41(3): p. 643–5.

52. Smith, K., et al., Rapid generation of fully human monoclonal antibodies specific to a vaccinating antigen. Nat Protoc, 2009. 4(3): p. 372–84.

53. Brooke, C.B., et al., Most influenza a virions fail to express at least one essential viral protein. Journal of virology, 2013. 87(6): p. 3155–62.

54. Wan, H., et al., Structural characterization of a protective epitope spanning A(H1N1)pdm09 influenza virus neuraminidase monomers. Nat Commun, 2015. 6: p. 6114.

55. Chen, Z., et al., Generation of live attenuated novel influenza virus A/California/7/09 (H1N1) vaccines with high yield in embryonated chicken eggs. J Virol, 2010. 84(1): p. 44–51.

56. Demaison, C., et al., High-level transduction and gene expression in hematopoietic repopulating cells using a human immunodeficiency [correction of imunodeficiency] virus type 1-based lentiviral vector containing an internal spleen focus forming virus promoter. Hum Gene Ther, 2002. 13(7): p. 803–13.

57. Martinez-Sobrido, L., et al., Hemagglutinin-pseudotyped green fluorescent protein-expressing influenza viruses for the detection of influenza virus neutralizing antibodies. J Virol, 2010. 84(4): p. 2157–63.

58. Reed, L.J. and H. Muench, A simple Method of Estimating Fifty Per Cent Endpoints. Am J Hyg, 1938. 27(3): p. 493–497.

59. Rowe, T., et al., Detection of antibody to avian influenza A (H5N1) virus in human serum by using a combination of serologic assays. J Clin Microbiol, 1999. 37(4): p. 937–43.

60. Lambre, C.R., et al., An enzyme-linked lectin assay for sialidase. Clin Chim Acta, 1991. 198(3): p. 183–93.

61. Couzens, L., et al., An optimized enzyme-linked lectin assay to measure influenza A virus neuraminidase inhibition antibody titers in human sera. J Virol Methods, 2014. 210C: p. 7–14.

62. Puleston, D.J., et al., Autophagy is a critical regulator of memory CD8(+) T cell formation. Elife, 2014. 3: p. e03706.

63. Southam, D.S., et al., Distribution of intranasal instillations in mice: effects of volume, time, body position, and anesthesia. Am J Physiol Lung Cell Mol Physiol, 2002. 282(4): p. L833–9.

64. Powell, T.J., et al., Pseudotyped Influenza A Virus as a Vaccine for the induction of Heterotypic Immunity. J Virol, 2012. 86(24): p. 13397–13406.

65. Joseph, T., et al., Evaluation of replication and pathogenicity of avian influenza a H7 subtype viruses in a mouse model. J Virol, 2007. 81(19): p. 10558–66.

66. Air, G.M., Influenza neuraminidase. Influenza Other Respir Viruses, 2012. 6(4): p. 245–56.

67. Wohlbold, T.J., et al., Vaccination with adjuvanted recombinant neuraminidase induces broad heterologous, but not heterosubtypic, cross-protection against influenza virus infection in mice. MBio, 2015. 6(2): p. e02556.

68. Monto, A.S., et al., Antibody to Influenza Virus Neuraminidase: An Independent Correlate of Protection. J Infect Dis, 2015.

69. Webster, R.G. and W.G. Laver, Preparation and properties of antibody directed specifically against the neuraminidase of influenza virus. J Immunol, 1967. 99(1): p. 49–55.

70. Lin, Y.P., et al., Neuraminidase receptor binding variants of human influenza A(H3N2) viruses resulting from substitution of aspartic acid 151 in the catalytic site: a role in virus attachment? J Virol, 2010. 84(13): p. 6769–81.

71. Hooper, K.A. and J.D. Bloom, A mutant influenza virus that uses an N1 neuraminidase as the receptor-binding protein. J Virol, 2013. 87(23): p. 12531–40.

72. Chen, Z., et al., The 2009 pandemic H1N1 virus induces anti-neuraminidase (NA) antibodies that cross-react with the NA of H5N1 viruses in ferrets. Vaccine, 2012. 30(15): p. 2516–22.

73. Easterbrook, J.D., et al., Protection against a lethal H5N1 influenza challenge by intranasal immunization with virus-like particles containing 2009 pandemic H1N1 neuraminidase in mice. Virology, 2012. 432(1): p. 39–44.

74. Wan, H., et al., Molecular basis for broad neuraminidase immunity: conserved epitopes in seasonal and pandemic H1N1 as well as H5N1 influenza viruses. Journal of virology, 2013. 87(16): p. 9290–300.

75. Zhou, F., et al., A triclade DNA vaccine designed on the basis of a comprehensive serologic study elicits neutralizing antibody responses against all clades and subclades of highly pathogenic avian influenza H5N1 viruses. J Virol, 2012. 86(12): p. 6970–8.

76. Schwartzman, L.M., et al., An Intranasal Virus-Like Particle Vaccine Broadly Protects Mice from Multiple Subtypes of Influenza A Virus. MBio, 2015. 6(4): p. e01044.

77. Harris, K., et al., Intramuscular immunization of mice with live influenza virus is more immunogenic and offers greater protection than immunization with inactivated virus. Virol J, 2011. 8: p. 251.

